# RasGRP1 promotes efficient intestinal tuft cell differentiation

**DOI:** 10.64898/2026.09.30.755407

**Authors:** Lauren A. Shechtman, Philippe Depeille, Eric Smith, Hong-Erh Liang, Tammie S. Y. Tam, Annabelle Duflock, Victor S. Cortez, Claire E. O’Leary, Alan Shen, Wouter Bos, Gabriela Fradiadakis, Alexis Combes, Hugo Gonzalez, Richard M. Locksley, Jeroen P. Roose

## Abstract

Intestinal stem and progenitor cells balance proliferation and lineage commitment to maintain epithelial homeostasis and adapt to environmental challenges. Tuft cells are a rare chemosensory lineage that rapidly expands in response to the cytokine IL-13 during helminth infection, yet how epithelial progenitor states influence this response remains incompletely understood. Using genetic mouse models, intestinal organoids, and single-cell approaches, we identify the Ras guanine nucleotide exchange factor RasGRP1 as an epithelial-intrinsic promoter of efficient tuft cell differentiation. *Rasgrp1* is expressed in rare crypt epithelial cells distinct from Lgr5-positive stem cells. Once cells entered the tuft cell differentiation trajectory, *Rasgrp1* deficiency has little impact on tuft cell maturation, including expression of the *Pou2f3* tuft cell transcription factor. Instead, RasGRP1 acts at the progenitor stage, where it restrains EGFR-driven proliferation and supports a transcriptionally heterogeneous transit-amplifying state, including a subset of cells expressing *Pou2f3*, that is largely lost upon *Rasgrp1* deficiency. Consistent with this model, EGFR inhibition restores tuft cell numbers in *Rasgrp1*-deficient organoids and cooperates with IL-13 to enhance tuft cell differentiation. *Rasgrp1*-deficient mice exhibit delayed tuft cell hyperplasia and inefficient helminth clearance. Conversely, constitutive epithelial RasGRP1 overexpression, uncoupled from its endogenous regulation, drives spontaneous tuft cell hyperplasia and accelerates parasite clearance, but alters the balance of intestinal epithelial lineages. Together, our findings identify RasGRP1 as a cell-intrinsic regulator that restrains progenitor proliferation and promotes responsiveness to IL-13, thereby coupling growth factor signaling, cell cycle control, and immune-driven epithelial lineage specification.

## Main Text

Differentiated epithelial cells in the small intestine (SI) are continuously replenished (∼3-5 days) by intestinal stem cells (ISCs) residing in the crypt niche (*1, 2*). ISCs undergo self-renewal and produce transit-amplifying (TAs) progenitor cells, which subsequently proliferate and differentiate into mature lineages with secretory (Paneth, Goblet, Enteroendocrine, Tuft) or absorptive (Enterocyte) functions (*2*). To maintain proper tissue homeostasis, ISCs and TAs must tightly balance proliferation and differentiation into appropriate lineages by integrating cell-intrinsic traits and extrinsic cues, such as BMPs-, EGF-, or immune-stimuli (*2–4*). TA progenitor cells have been proposed to modulate the composition of differentiated cell types (*5*), However, the regulatory mechanisms that ensure progenitor cells make the desired ratio of binary proliferation- and differentiation-fate choices across diverse environmental contexts are unclear. Further, the lack of defined discriminatory or functional markers in the TA progenitor population has hampered discovery in this area.

Rare chemosensory tuft cells in the SI detect luminal parasites and release the cytokine interleukin-25 (IL-25), thereby activating type II innate lymphoid cells (ILC2s) and type 2 helper T (Th2) cells to secrete IL-13 (*6–8*). This extrinsic IL-13 cue drives tuft and goblet cell production from undifferentiated epithelial cells, a feed-forward circuit that continues as infection persists (*6–9*). The resulting ‘*weep & sweep*’ response, characterized by increased mucus production and enhanced smooth muscle activity, facilitates worm expulsion (*9–11*). Recent organoid studies revealed that IL-13, mimicking infection *in vitro*, may also drive a regenerative response and steer tuft cells back into the cell cycle and a multipotent state (*12*). Collectively, these findings imply that IL-13-mediated fate choices are modulated by unknown intrinsic features of epithelial cells.

Our previous work described how RasGRP1 (Ras guanyl nucleotide releasing protein 1), a Ras guanine nucleotide exchange factor (RasGEF), dampens EGF-driven proliferation in the mouse SI and in patients and mouse models with colorectal cancer (*13*). Using different genetic mouse models, organoids, and single cell resolution techniques that enabled detailed interrogation of rare TA and tuft cells, we show here that RasGRP1 marks a rare TA progenitor subpopulation and is a critical driver of the tuft cell lineage and functions by dampening cell-intrinsic cell cycle activity, thereby licensing tuft cell specification through IL-13.

### *Rasgrp1* deficiency results in decreased tuft cell generation

Based on low-resolution IHC, we previously reported that RasGRP1 protein is expressed in an undefined region of SI crypts (*13*). To precisely define its spatial distribution, we performed RNAscope and high-resolution imaging on proximal SI tissue sections from WT mice and detected *Rasgrp1* transcripts in rare epithelial cells within the crypt compartment where TAs reside; an area distinct from the crypt region with *Lgr5*-positive stem cells (Fig. 1A, arrows). Since RasGRP1 dampens EGF-driven proliferation (*13*) and TA zone proliferation characteristics have been suggested to impact composition of differentiated cell types (*5*), we postulated that RasGRP1 may regulate lineage commitment in the SI.

**Fig. 1.**
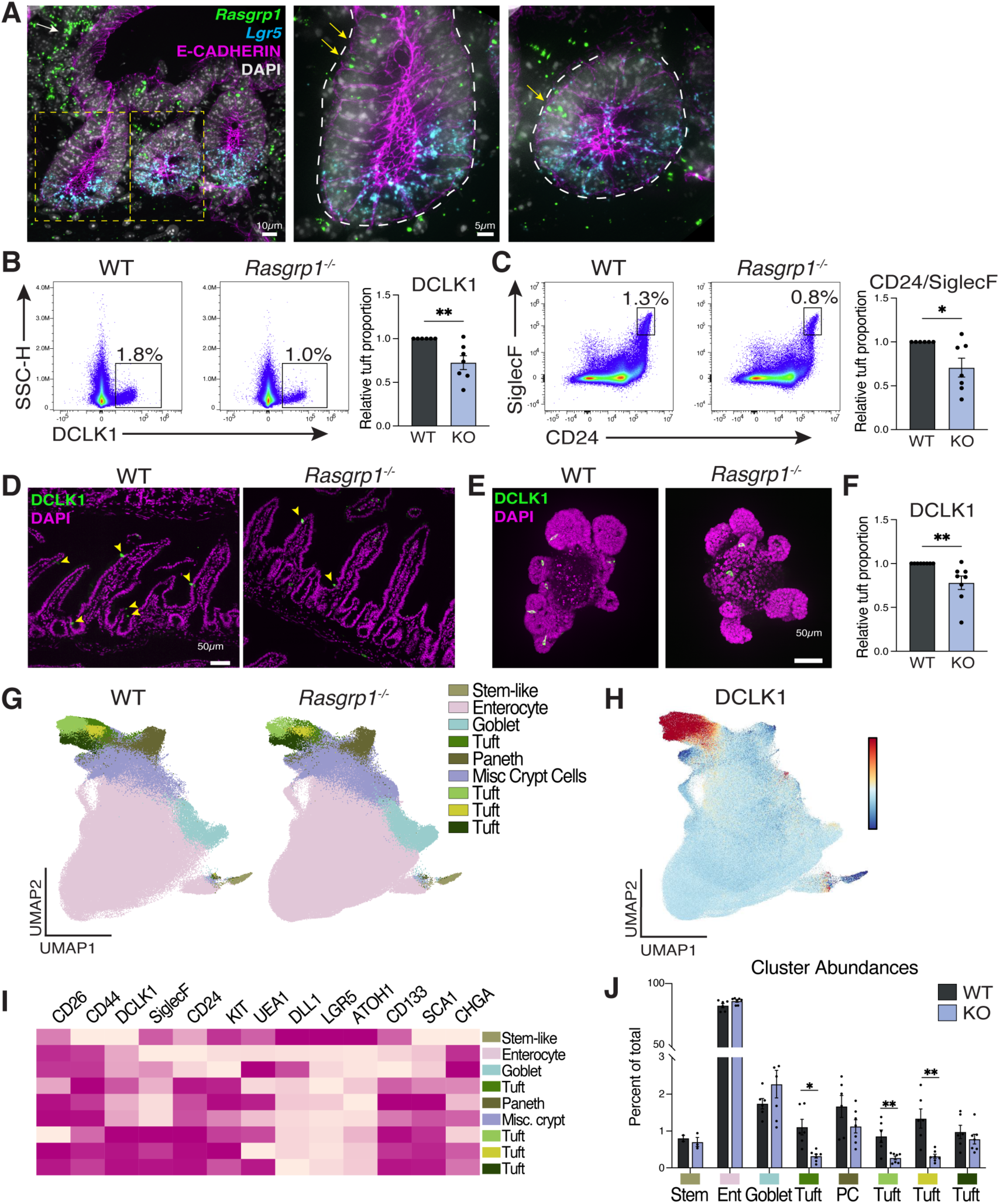
RasGRP1 is necessary for efficient tuft cell generation. (**A**) *Rasgrp1* mRNA expression (green, arrows) in rare E-CADHERIN^+^ (magenta) epithelial cells, and lamina propria (white arrow) in proximal SI crypt (white outline), exclusive from *Lgr5* mRNA expression (cyan). Nuclei stained with DAPI (white). Scale bar, 10µm and 5µm as indicated. (**B, C**) Representative flow cytometry and quantification of DCLK1^+^ and CD24^+^/SiglecF^+^ tuft cells in WT (n=6) and *Rasgrp1^−/−^*(n=7) duodenum. Tuft cell proportions quantified relative to total EPCAM^+^CD45^low/neg^ cells. (**D**) Representative IF images of tuft cells (DCLK1, green, arrowheads) in WT and *Rasgrp1^−/−^*duodenum. Nuclei stained with DAPI (magenta). Scale bar, 50µm. (**E, F**) IF and quantified flow cytometry of DCLK1^+^ tuft cells in WT and *Rasgrp1^−/−^*duodenal organoids. Tuft cell proportions quantified relative to total EPCAM^+^CD45^low/neg^KI67^−^ CASPASE^−^ differentiated epithelial cells. Nuclei stained with DAPI (magenta). Scale bar, 50µm. (**G**) UMAP visualization colored by differentiated cell types of spectral flow cytometry data generated from WT and *Rasgrp1^−/−^*crypts. UMAP represents all EPCAM^+^CD45^low/neg^KI67^−^ CASPASE^−^ differentiated epithelial cells. (**H**) UMAP visualization of DCLK1 protein expression in merged WT and *Rasgrp1^−/−^*UMAP. (**I**) Heatmap of normalized marker expression in each cluster from (**G**). (**J**) Abundance of each cell type cluster in WT and *Rasgrp1^−/−^*datasets. Note that frequencies of *Rasgrp1^−/−^*goblet cells and enterocytes were not significantly increased with this analysis method Bars depict mean ± standard error of the mean (SEM). Each data point is a biological replicate (1 mouse or 2 pooled organoid wells). *p<0.05, **p<0.01 (unpaired two-tailed t-tests). Data are representative of at least three experiments (B, C, F, and J).

To test this, we established a multiparameter spectral flow cytometry panel (Table S1) that allows us to resolve the major epithelial cell types in the murine SI and in derived organoids (Fig. S1A-E, S2A). We first applied manual gating on spectral flow cytometry data within the EPCAM^+^CD45^low/neg^ population to quantitatively assess known differentiated epithelial cell types (Fig. S1B). Such manual gating of stained single cells from crypts of cohoused WT and *Rasgrp1* knockout (*Rasgrp1^−/−^*, KO) duodenum revealed that KO tissue contained fewer tuft cells, compared to WT controls, using either DCLK1 or CD24/SiglecF as markers for this lineage (Fig. 1B,C and Figs. S2B, C), a finding we confirmed using DCLK1 immunofluorescence (IF) (Fig. 1D). Manual gating strategies, relying on other stereotypic markers as proxy for different lineages, also revealed increased frequencies of goblet cells and enterocytes in KO SI, whereas Paneth- and enteroendocrine-(EEC) cell frequencies were unchanged (Fig. S2D).

To determine what lineage differentiation defects observed in KO mice are purely epithelial-intrinsic and retain in the stem/progenitor compartment, we next generated SI organoids from WT and KO mice (Fig. S2A) and analyzed late passages (P3–8), which are largely devoid of immune and stromal cells and retain the major epithelial lineages *ex vivo* (*14*) Tuft cells were the unique cell population reduced in KO organoids at steady state and grown in conventional organoid media (Figs. 1E, F, S2E), indicating that RasGRP1 has a previously unrecognized epithelial-intrinsic function, promoting tuft cell specification, likely at an early stage of cell specification.

We next applied Uniform Manifold Approximation and Projection (UMAP) dimensionality reduction analysis to explore cell types and global data structure of single cells from crypts of WT and *Rasgrp1* knockout mice. These analyses capitalize on the cumulative discriminatory power of the extensive antibody panel and provided an unbiased overview of most epithelial populations, except for rare EECs and specific stem/progenitor subtypes (Figs. S1C, D) and were highly reproducible across biological replicates (Fig. S1E). To computationally resolve heterogenous cell subsets in an unbiased manner, we next plotted EPCAM⁺CD45^low/neg^KI67^−^ CASPASE^−^ cells to assess the presence of subsets of differentiated cells (Figs. 1G). We identified four tuft cell clusters, all expressing high levels of DCLK1 (Fig. 1H) but showing varying expression levels of other tuft cell markers, such as KIT, SiglecF, and UEA1 (Fig. 1I). Furthermore, three of the four tuft clusters were significantly reduced in KO SI, while other differentiated lineages (i.e., goblet cells, enterocytes) did not display significant changes with this analysis method (Fig. 1J). Given the unique expression pattern of RasGRP1 in a subset of progenitor cells and the specific reduction of tuft cells as the only affected differentiated population in KO organoids, we focused on defining the role of RasGRP1 in regulating the tuft cell lineage in this study.

### Immature tuft cell subset production is regulated by RasGRP1

The transcription factor POU2F3 is crucial for tuft cell development, as mice lacking *Pou2f3* are completely devoid of tuft cells (*15*). During infection, tuft cells sense helminth-derived metabolites and release IL-25, which activates ILC2s to secrete IL-13 and engage the tuft cell circuit. Accordingly, mice lacking *Stat6* (Signal Transduced and Activation of Transcription 6), a transcription factor that drives IL-13 responsive genes, are unable to expand tuft and goblet cell numbers during helminth infection or clear worms (*8, 16*). None of our previous studies on RasGRP1 (*13, 17–23*) suggested a role in IL-13/Stat6 signaling. We therefore hypothesized that *Rasgrp1* deficiency impacts tuft cell development in a manner distinct from the extrinsic IL-13 cue and performed single-cell RNA sequencing (scRNA-seq) to interrogate this idea.

*Rasgrp1^−/−^*SI exhibited reduced numbers of SiglecF^+^/CD24^+^ tuft cells, but the levels of expression for these two cell surface markers were comparable between WT and KO SI (Fig. 1C); thus, we used SiglecF and CD24 (*24*) to enrich for tuft cells and generate high-resolution gene expression datasets from this rare cell population. Unsupervised clustering partitioned cells into 10 epithelial populations, annotated using established lineage markers of intestinal cell types (Fig. 2A-C, S3A-D) (*25*). Five clusters expressed tuft cell markers, including *Cd24a*, *Siglecf*, *Dclk1*, *Pou2f3*, and *Trpm5* (Fig. 2B, C) (*15, 25*). Our intentionally permissive sorting on SiglecF and CD24 also captured some cells in two goblet cell clusters (*Muc2*^+^, *Tff3*^+^ Fig 2A, C, orange), two enterocyte clusters (*Fabp2*^+^, *Alpi*^+^, Fig 2A, C, blue), and one transit amplifying (TA) cluster (*Dmbt1*^+^, *Mki67*^+^, Fig 2A, C, green). Differential gene expression analysis of WT cells further confirmed the identity of these 10 clusters, with goblet cells expressing *Defa3* and *Atoh1*, enterocyte clusters expressing *Reg3a* and *Fabp2*, and TA clusters *Ccnb1,* and *Melk* (Fig. S3C, Table S2). Goblet cell cluster fractions were higher in the KO dataset, while tuft cell clusters were lower (Fig. 2D), resembling our spectral flow data. Notably, the TA cell fraction was modestly increased in KO, compared to WT, indicating that *Rasgrp1* loss did not cause a general TA progenitor cell defect, at least from a numeric perspective Fig. 2D).

**Fig 2.**
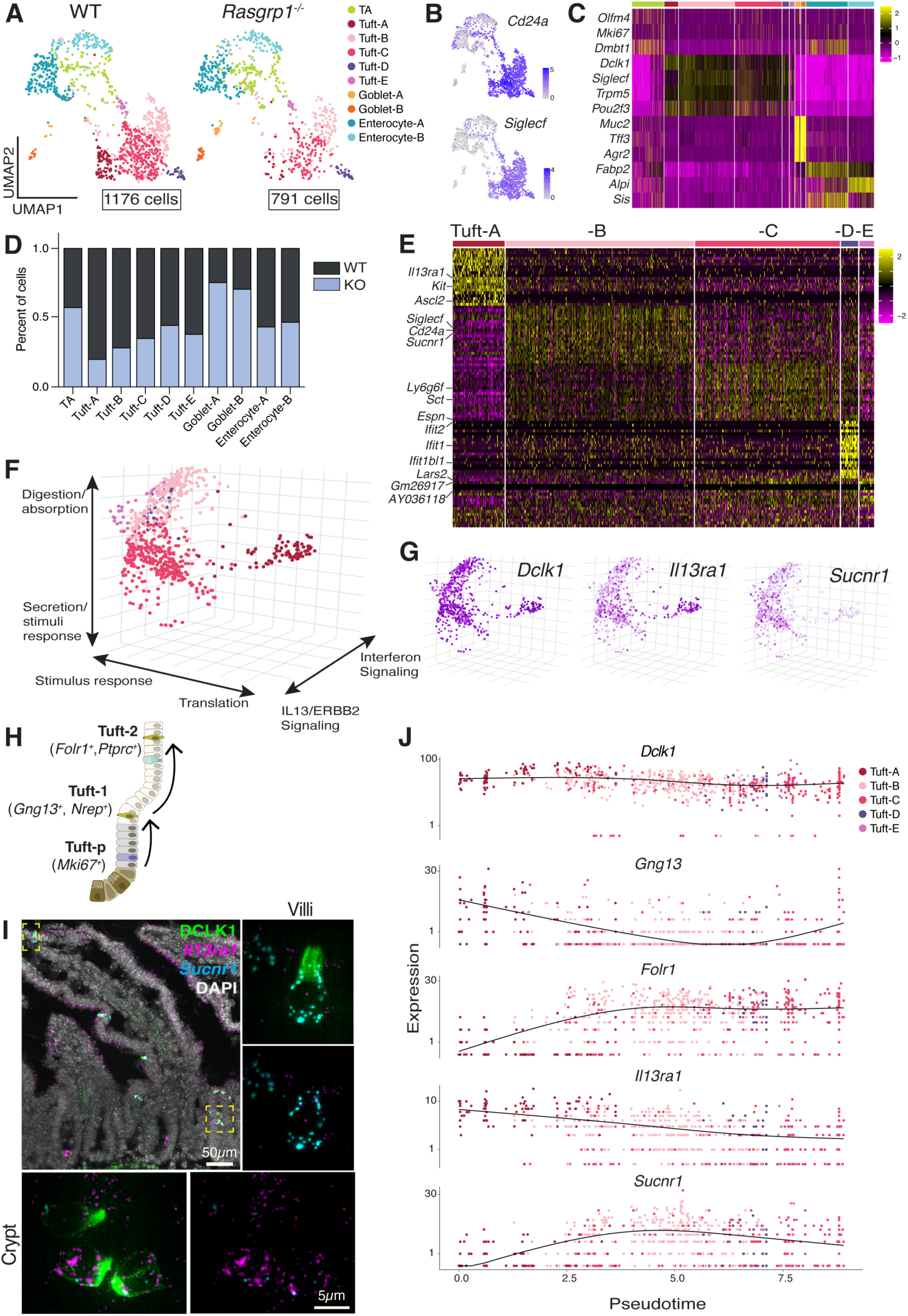
RasGRP1 is a regulator of immature tuft cell production. (**A**) UMAP visualization of EPCAM^+^CD45^neg/low^CD24^+^SiglecF^+^ cells isolated from WT (n=3) and *Rasgrp1^−/−^*(n=3) intestines. Cells are colored by annotated cell subsets. (**B**) Feature plots of tuft markers, *Cd24a,* and *Siglecf*. (**C**) Heatmap of canonical cell type gene expression across clusters in merged dataset. (**D**) Quantified distribution of clusters in WT and *Rasgrp1^−/−^*datasets. (**E**) Heatmap of the top 20 differentially expressed genes across tuft cell clusters in WT dataset. (**F**) Diversity of tuft cell clusters in WT cells evaluated by diffusion map analysis. (**G**) Normalized expression of *Dclk1, Il13ra1,* and *Sucnr1* across diffusion map. (**H**) Schematic illustrating published tuft cell subsets. (**I**) Representative RNAscope image of *Il13ra1* (magenta) and *Sucnr1* (cyan) mRNA expression in crypt and villi tuft cells (DCLK1, green). Nuclei stained with DAPI (gray). Scale bar, 50µm and 5µm. (**J**) Expression trajectories of tuft cell subset genes in WT tuft cell clusters over pseudotime.

To better understand the progressive stages of normal tuft cell development through, we first performed differential expression exclusively on WT clusters and subjected the WT dataset to diffusion component analysis, a non-linear dimensionally reduction technique which captures geometric structure and functional states in scRNA-seq data (*26–29*). Both methods identified tuft-A as an immature state enriched in mRNA translation, a feature that has been associated with transitional cell states (*30, 31*), and marked by high expression of *Il13ra1*, *Ascl2*, *Cd44*, and *Kit* (Fig. 2E-G, Table S2). In contrast, tuft clusters B-E displayed signatures of more mature, specialized tuft cells (Figs. 2E-G). Tuft-B highly expressed canonical tuft cell genes (*Sucnr1*, *Siglecf*, and *Cd24a*), linked to succinate sensing and inflammatory responses (*32–34*). Diffusion analysis further linked this cluster to digestion and absorption pathways. Tuft-C was enriched for *Sct, Espn,* and *Ly6g6f* as well as *Neurod1, Clca2,* and *Folr1*, which contain strong associations to secretory and stimulus-response functions. Tuft-D expressed interferon-stimulated genes (*Ifit1, Ifit2, Ifit1bl1*), consistent with tuft cell-intrinsic antiviral functions (*35–37*). Given the small number of cells in the Tuft-E cluster, we refrained from proposing function to Tuft-E based on transcriptional characteristics. In sum, our differential expression and 3D diffusion map analysis demonstrated that tuft-A cells represent an immature state, while tuft clusters B-D constitute mature subtypes with distinct functional specializations ranging from nutrient sensing to antiviral defense (Fig. 2F).

Prior single-cell studies have identified two major tuft cell subsets, tuft-1 and tuft-2 (*25, 38*), and recently described tuft progenitor signature (Tuft-p, Fig. 2H) (*39*). To create unity, we therefore further analyzed our tuft-A-E dataset in the context of this framework. Collectively, tuft-A and tuft-1 signatures strongly overlap, whereas tuft clusters B-E align with tuft-2 (Fig. S3E-H), reinforcing the concept that tuft-1 corresponds to an immature state located within intestinal crypts, and tuft-2 to more differentiated functional programs in the villi (Fig. 2F, H) (*38*). The recent described tuft progenitor signature (Tuft-p, Fig. 2H) mapped closest to our enterocyte and TA clusters, suggesting that these populations may serve as a common tuft progenitor (Fig. S3H). We did not observe similarities to tuft-3 and tuft-4, albeit that injury conditions were not assessed in our study. (*12*). Lastly, in agreement with our *Rasgrp1* RNAscope data in very rare progenitor cells (Fig. 1A) we could not detect *Rasgrp1* transcripts in our own data set here, nor in any of the publicly available datasets (*12, 25, 39, 40*).

We noted enriched expression of the IL-13 receptor (*Il13ra1*) in tuft-A, whereas the succinate receptor (*Sucnr1*) was broadly expressed in differentiated epithelial cells but enriched in tuft-B (Fig. 2E, G). Orthogonal RNAscope approaches on proximal WT SI tissue sections confirmed *Il13ra1* expression is highly enriched in crypt-localized DCLK1-positive cells with little but present expression in villus tuft cells, and throughout the intestinal epithelium, whereas *Sucnr1* was restricted to villus tuft cells (Fig. 2I). Our relatively high number of single tuft cell datapoints allowed for monocle pseudotime analysis (*41*) (Fig. S4A), to assess the proposed model of tuft cell lineage dynamics (Fig. 2H). While *Dclk1*, and *Pou2f3* expression remained constant (Fig. 2J, S4B), tuft-1/tuft-A marker expression (*Gng13* and *Il13ra1*) progressively decreased over pseudotime, and tuft-2/tuft-B-E markers (*Folr1*, *Sucnr1*) increased (Fig. 2J). Taken together, our data analyses of WT SI support the model of zonated tuft cell populations and refines the characterization of distinct tuft cell subsets within a continuum of cell states.

### RasGRP1 restricts EGFR-driven proliferation to license IL-13 cues

*Rasgrp1* deficiency profoundly impacted the immature tuft compartment, as tuft-1/tuft-A was reduced by ∼80% of WT cell numbers, whereas tuft clusters B-E were reduced by 55-70% (Fig. 2D). Since *Rasgrp1*-deficient mice particularly lack this *Il13ra1-*expressing tuft-1/tuft-A subset (Fig. S4C), we next infected WT and KO mice with the helminth *Nippostrongylus brasiliensis* (*Nippo*) and assessed worm clearance and tuft cell hyperplasia, 7- and 9-days post infection (Fig. 3A). KO mice exhibited impaired worm clearance with detectable worms at day 9 when WT mice had cleared infection (Fig. 3B). Consistent with this result, tuft cell hyperplasia was delayed in KO at day 7, as measured by spectral flow (Fig. 3C) and IF on SI sections (Fig. 3D). Intriguingly, tuft cell numbers in KO SI approached WT levels by day 9, likely due to persistent signals from helminths lingering in the KO SI lumen at day 7 (Fig. 3C).

**Fig. 3.**
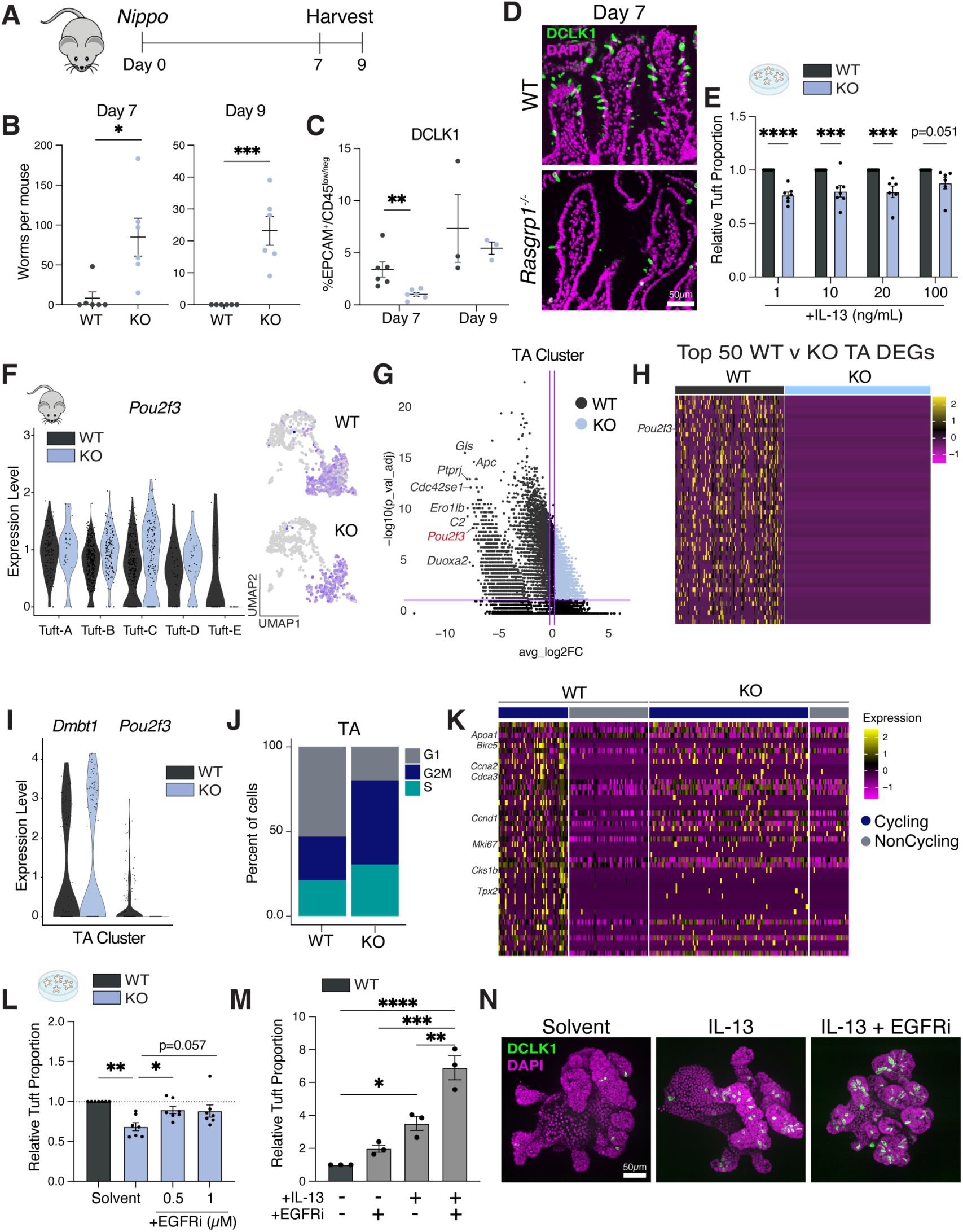
RasGRP1 controls tuft cell generation and hyperplasia through regulation of the cell cycle. (**A**) Experimental diagram for *N. brasilienis* infection. (**B**) Worm counts in mice infected 7, and 9 days with *N. brasilienis.* (**C**) Flow cytometry quantification of DCLK1^+^ tuft cells in mice infected with *N. brasilienis* for 7, and 9 days. (**D**) Representative IF images of tuft cells (DCLK1, green) in WT and *Rasgrp1^−/−^*duodenum, 7 days post infection. Scale = 50µm. (**E**) Quantified flow cytometry of DCLK1^+^ tuft cells in WT duodenal organoids with increasing concentrations of IL-13. (**F**) Violin plot and UMAP visualization showing expression of *Pou2f3* in indicated WT and *Rasgrp1^−/−^*clusters. (**G**) Volcano plot showing genes differentially expressed in WT versus KO TA cells. Each point represents a gene, with color indicating direction of change (up in KO vs up in WT). Horizontal and vertical purple lines indicate thresholds for significance (adjusted p-value < 0.05) and log2 fold change > 0.2, respectively. The top 8 up-and down-regulated genes are labeled. (**H**) Heatmap of top 50 genes differentially expressed in WT versus KO TA cells, as in **G**. (**I**) Violin plot and UMAP visualization showing expression of *Dmbt1* and *Pou2f3* in the TA cluster from WT and *Rasgrp1^−/−^*datasets. (**J**) Percentage of cells in each cell cycle phase in WT and *Rasgrp1^−/−^*TA clusters. (**K**) Heatmap showing genes significantly up or down-regulated in cycling (G2M/S) and non-cycling (G1) TA cells in WT and KO datasets. (**L**) Quantified flow cytometry of DCLK1^+^ tuft cells in *Rasgrp1^−/−^*duodenal organoids treated with increasing concentrations of gefitinib (EGFRi), relative to WT. (**M**) Quantified flow cytometry, and (**N**) representative IF images of DCLK1^+^ tuft cells in WT duodenal organoids treated with 20ng/mL IL-13, with or without the addition of 1µM gefitinib (EGFRi). In **E**, **L**, and **M**, tuft cell proportions quantified relative to total EPCAM^+^CD45^low/neg^KI67^−^ CASPASE^−^ differentiated epithelial cells. Bars depict mean ± standard error of the mean (SEM). Each data point is a biological replicate (1 mouse or 2 pooled organoid wells). *p<0.05, **p<0.01, ***p<0.001, and ****p<0.0001 (unpaired two-tailed t-tests for B, 1way ANOVA for E, L, M and 2way ANOVA for C). Data are representative of at least two (B, C, M) or three (E, L) experiments.

Since our KO SI organoids (Figs. 1E, F) retained the steady-state tuft lineage specification defect seen *in vivo*, we next mimicked worm infection *in vitro* by exposing WT and KO organoids grown in conventional organoid media to IL-13 (*6, 42, 43*). We tracked tuft cell hyperplasia as a function of increasing concentrations of IL-13 (Fig. S4D) and made two observations. First, while IL-13 induced tuft cell expansion in both WT and KO organoids by day 5 of culture (Fig. S4E), KO organoids consistently generated fewer tuft cells than WT controls (Fig. 3E). Secondly, in both WT and KO organoids tuft cell hyperplasia demonstrated a plateau in the response to increasing IL-13, as reported previously (*39*). Combining the data of *Nippo* infection *in vivo* and IL-13 stimulation on organoids, we concluded that RasGRP1 plays an epithelial-intrinsic role, somehow optimizing efficient IL-13-driven tuft cell hyperplasia and helminth clearance.

To understand mechanistically how RasGRP1 may regulate production of tuft-1/tuft-A cells from epithelial cells, we first analyzed several genes previously implicated in tuft cell differentiation (*15, 44, 45*). Expression of lineage-defining markers such as *Dclk1, Sox4, Cdc42,* and the transcription factor *Pou2f3*, appeared largely normal across tuft A-E populations in *Rasgrp1* deficient animals (Fig. 3F, S4F-H), arguing that RasGRP1 does not control tuft cell lineage identity once specification has occurred. In concordance, KO tuft cell clusters exhibited relatively few transcriptional changes (maximum of 378 DEGs; Table S2). By contrast, differential expression analysis of the TA compartment from WT and KO SI identified 6,227 DEGs. Whereas WT TA cells displayed a broad transcriptional program with distinct cell-to-cell variation, these varied single cell transcriptional states were almost entirely lacking when *Rasgrp1* is absent (Figs. 3G, H, S4I, Table S2). Notably, a portion of *Dmbt1*-positive WT TA cells express low and varied levels of *Pou2f3* (Fig. 4I), a feature that was absent in *Rasgrp1* deficient *Dmbt1*-positive counterparts (Fig. 4I). These results indicate that RasGRP1 functions to support a poised state of TA progenitors.

**Fig. 4.**
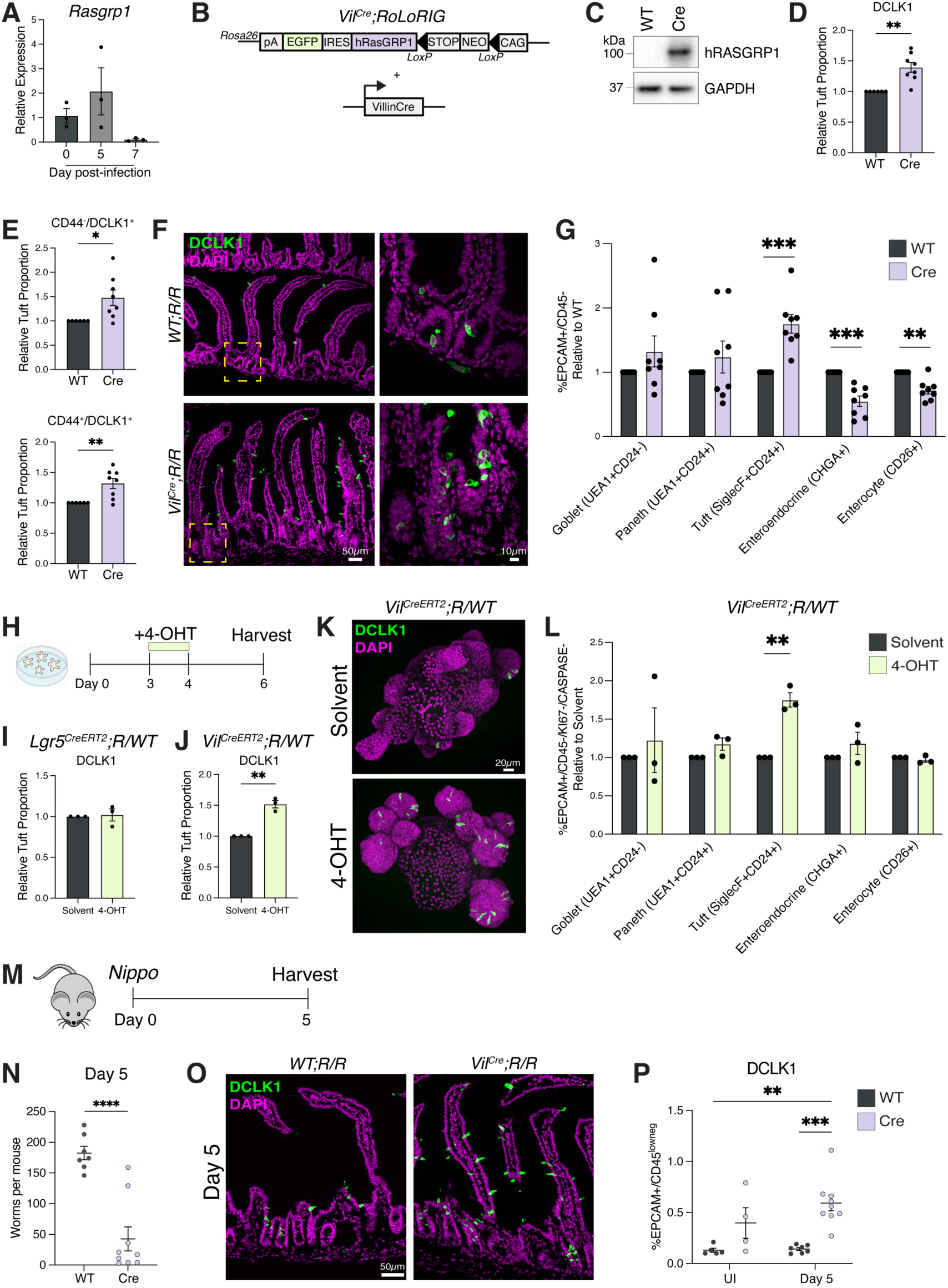
Elevated RasGRP1 expression in epithelial cells drives tuft cell generation and hyperplasia. (**A**) Transcript analysis of *Rasgrp1* in sorted duodenal epithelial cells of WT uninfected (day 0), and *Nippo* infected (day 5, 7) mice (n=3). (**B**) Schematic of *RoLoRIG* mouse construct. (**C**) Western blot showing levels of indicated proteins in duodenum of *R/R;WT* and *Vil^Cre^;R/R* mice. (**D**) Quantified spectral flow cytometry data of EPCAM^+^CD45^low/neg^ tuft cells isolated from *R/R;WT* (n=6) and *Vil^Cre^;R/R* (n=8) duodenum. (**E**) Quantified EPCAM^+^CD45^low/neg^ crypt (CD44^+^DCLK1^+^) and villi (CD44^−^ DCLK1^+^) tuft cell proportions in *R/R;WT* and *Vil^Cre^;R/R* duodenum. (**F**) Representative IF images of tuft cells (DCLK1, green) in *R/R;WT* and *Vil^Cre^;R/R* mice duodenum. Nuclei stained with DAPI (magenta). Scale = 50µm, and 5µm. (**G**) Quantified spectral flow cytometry data of differentiated EPCAM^+^CD45^low/neg^ cells isolated from *R/R;WT* (n=6) and *Vil^Cre^;R/R* (n=8) intestines. (**H**) Experimental diagram of 4-hydroxytamoxifen (4-OHT) treatment of *Lgr5^CreERT2^*;*R/WT* and *Vil^CreERT2^*;*R/WT* organoids. (**I, J**) Quantified spectral flow cytometry of DCLK1^+^ tuft cells in 4-OHT and solvent treated (**I**) *Lgr5^CreERT2^*;*R/WT* and (**J**) *Vil^CreERT2^*;*R/WT* duodenal organoids. Tuft cell proportions quantified relative to total EPCAM^+^CD45^low/neg^KI67^−^ CASPASE^−^ differentiated epithelial cells. (**K**) Representative IF images of tuft cells (DCLK1, green) in 4-OHT and solvent treated *Vil^CreERT2^*;*R/WT* organoids. Nuclei stained with DAPI (magenta). Scale bar, 20µm. (**L**) Quantified spectral flow cytometry data of differentiated EPCAM^+^CD45^low/neg^KI67^−^ CASPASE^−^ cells isolated in 4-OHT and solvent treated *Vil^CreERT2^*;*R/WT* organoids. (**M**) Experimental diagram for *N. brasilienis* infection. (**N**) Worm counts in mice infected 5 days with *N. brasilienis.* (**O**) Representative IF images of tuft cells (DCLK1, green) in *R/R;WT* and *Vil^Cre^;R/R* duodenum 5 days post *N. brasilienis* infection. Scale = 50µm. (**P**) Flow cytometry quantification of DCLK1^+^ tuft cells in mice infected with *N. brasilienis* for 5 days. Bars depict mean ± standard error of the mean (SEM). Each data point is a biological replicate (1 mouse or 2 pooled organoid wells). *p<0.05, **p<0.01, ***p<0.001, and ****p<0.0001 (unpaired two-tailed t-tests for D,E,G,I,J,L, and N and 1way ANOVA for A, and P). Data are representative of at least two (I, J, N, P) or three (D, E, G) experiments.

TA cells and their proliferation status have been proposed to influence the composition of differentiated cells (*5*). Our prior work in colorectal cancer cells demonstrated that RasGRP1 suppresses proliferative SOS-Ras-ERK signals, downstream of the Epidermal Growth Factor receptor (EGFR) (*13*). We therefore postulated that altered cell cycle dynamics in *Rasgrp1* deficient TA cells could affect their responsiveness to IL-13 and thereby contribute to the tuft cell defect in *Rasgrp1* KO mice. Of all duodenal cells analyzed by scRNAseq, ∼64% of KO cells were in G2M/S phase, compared to only ∼26% of WT cells (Fig. S4J, K). When examined by specific cluster, the TA cluster and tuft A-E clusters contained very high percentages of cells in G2M/S phase, relative to WT counterparts (Fig. 3J, S4J, K). To further dissect the relationship between cell cycle dynamics and gene expression programs, we performed differential expression analysis comparing cycling (G2M/S) versus non-cycling (G1) cells. In WT TA cells, there is a clear distinction in the gene programs between cycling TA cells, expressing *Mki67, Ccna2,* and *Ccnd1*, and non-cycling cells (Fig. 3K), indicating that cell cycle dynamics may regulate TA cell differentiation trajectories. By contrast, KO TA cells did not demonstrate such clear demarcation (Fig. 3K).

We postulated that dampening of proliferative EGFR signals in *Rasgrp1*-deficient progenitor cells may restore the poised state that favors tuft cell differentiation. To test this hypothesis, we first treated WT and KO SI organoids with the EGFR inhibitor (EGFRi) gefitinib, (*21*) a condition previously shown to promote cell cycle exit (*46–48*). KO organoids treated with 0.5 or 1µM gefitinib in conventional organoid media restored steady-state tuft cells numbers, comparable to WT (Fig. 3L). Further, combination of EGFRi and IL-13 enhanced tuft cell hyperplasia in organoids, beyond levels seen with either treatment alone (Fig. 3M, N). Together, these findings demonstrate that RasGRP1 enables a poised state so that a subset of TA progenitor cells express low levels of *Pou2f3*, supporting steady state tuft cell lineage commitment. Further, RasGRP1 restrains epithelial cell-intrinsic proliferation thereby licensing of cell-extrinsic IL-13 cues, enabling efficient but controlled tuft hyperplasia responses.

### RasGRP1 overexpression drives spontaneous tuft cell hyperplasia and worm clearance

Given the narrow expression pattern of *Rasgrp1* in the crypt at steady state (Fig. 1A), this raised the question of how RasGRP1 promotes efficient tuft cell hyperplasia during parasite infection. To explore this, we sorted epithelial cells from SI uninfected (day 0), and at early (day 5), and late (day 7) stages of *Nippo* infection. qPCR analysis revealed an initial upward trend of *Rasgrp1* expression, followed by a collapse by day 7, coinciding with peak tuft cell hyperplasia (Fig. 4A). These data suggest that *Rasgrp1* expression is a dynamically regulated, epithelial-intrinsic licensing factor, driving tuft cell specification.

To test our model wherein RasGRP1 functions as a gatekeeper in TA cells, biasing towards the tuft lineage while protecting sufficient generation of other epithelial cell subsets, we used a Rosa26 knock-in line (*RoLoRiG*, *RR*) carrying a CAG-driven LoxP-STOP-LoxP-hRASGRP1-IRES-EGFP cassette (Fig. 4B) (*22*). Human RasGRP1 shares 92-95% amino acid similarity with the murine protein and its expression is tolerized in mice (*19, 22*). Crossing of *RR* mice with *Villin-Cre* (*Vil^Cre^*) mice resulted in specific expression of human RASGRP1 in intestinal epithelium of *Vil^Cre^;R/R* animals (homozygous *RR*, *Villin-Cre^+/−^*), detected with our antibody, specific to human RASGRP1 (Fig. 4B, C, S5A).

Proximal SI from uninfected *R/R;VCre* mice revealed spontaneous hyperplasia of DCLK1^+^ tuft cells assessed by spectral flow cytometry, compared to littermate controls (Figs. 4D). a marker previously shown to distinguish crypt and villi cells (*40, 49*), we found that both crypt- and villus-tuft cell proportions were elevated in *Vil^Cre^;R/R* mice (Figs. 4E, S5B). IF approaches revealed striking patterns of DCLK1 expression in the crypts of *Vil^Cre^;R/R* animals at steady state (Fig. 4F), indicating that elevated RasGRP1 can drive tuft cell production across the crypt-villus axis. Spectral flow analysis of EPCAM⁺CD45^low/neg^ differentiated cells from *Vil^Cre^;R/R* SI and controls demonstrated that RasGRP1-driven tuft cell hyperplasia comes at a consequence of other cell lineages. Both enterocytes but also EEC proportions were reduced upon constitutive overexpression of RasGRP1 from the *Villin* promoter. (Fig. 4G).

Our RNAscope data indicates that *Rasgrp1* is expressed in a very rare cell type that is different from *Lgr5*-positive stem cells (Fig 1A). We therefore genetically explored through what crypt cell subset RasGRP1 can drive tuft cell hyperplasia. To distinguish whether RasGRP1 overexpression acts acutely within TA cells rather than ISCs, we employed two inducible Cre systems. *RoLoRiG* mice were crossed with *Lgr5^CreERT2^*, and *Villin^CreERT2^* lines to generate *Lgr5^CreERT2^;R/WT* and *Vil^CreERT2^;R/WT* animals, respectively, each heterozygous for both the *RoLoRiG* and *Cre* alleles. Duodenal epithelial organoids derived from these mice were treated with 4-hydroxytamoxifen (4-OHT) to induce recombination as previously described (*50–52*) (Fig. 4H). *Lgr5^CreERT2^;R/WT* organoids treated for 24hrs with 4-OHT and assayed 2 days later exhibited no change in the proportion of tuft cells or other epithelial lineages relative to solvent-treated controls (Figs. 4I, S5C). By contrast, *Vil^CreERT2^*;*R/WT* organoids resulted in elevated tuft cell numbers in this set up, as assessed by flow cytometry and IF (Figs. 4J-L). Lastly, like the unique tuft cell defect in *Rasgrp1*-deficient SI organoids (Fig. S2C), the acute and short-term induced RASGRP1 overexpression in *Vil^CreERT2^*;*R/WT* organoids exclusively enhanced tuft cell specification, without affecting other differentiated cell lineages (Fig. 4L).

Finally, to compare the *in vivo* consequence of epithelial-intrinsic overexpression of RasGRP1 to the WT situation of dynamically regulated RasGRP1 expression in the context of altered cell-extrinsic cues, we challenged our *Vil^Cre^;R/R* model and control mice with *Nippo.* Here, we purposely analyzed the duodenum 5 days post infection to test our hypothesis that RasGRP1 overexpression may enhance the ‘*weep & sweep*’ response (Fig. 4M). At day 5, control animals displayed 150-200 worms per mouse and the tuft cell hyperplasia program had not yet engaged to the level that is seen at day 7 (Figs. 4N-P, and 3B). By contrast, *Vil^Cre^;R/R* mice harbored significantly fewer worms in the duodenum at day 5 and the elevated steady-state tuft cell numbers are not further increased, presumably because of the lack helminth-derived metabolites (Figs. 4N-P). Collectively, these data demonstrate the need for dynamically controlled RasGRP1 expression in TA progenitor cells so that balanced lineage differentiation can occur in a coordinated response to external cues in the intestine.

## Discussion

In the small intestine, rare tuft cells share sparsity with enteroendocrine cells (EEC), indicating tightly controlled commitment towards these lineages. Both specialized, secretory cell lineages display lineage development as a continuum, as described in previous studies (*12, 39, 53*) and our tuft A-E subsets described here. The tuft cell lineage demonstrates a 10-fold expansion upon helminth infection, driven by extrinsic IL-13 (*6–8*); however, this IL-13 cue is limited in its capacity to induce tuft cell hyperplasia (*39, 43*). We discovered that *Rasgrp1* is expressed in rare crypt cells in a zone that is distinct from *Lgr5*-expression ISCs and is necessary for proper tuft cell production during homeostasis, and hyperplasia upon helminth infection. Further, *Rasgrp1* expression is dynamic, induced upon *Nippo* infection, and declining at time points when worms are cleared. Here, we discovered that the tuft lineage is under control of combinatorial extrinsic and intrinsic cues. In essence, tuft cell commitment requires two critical inputs: (a) the cell-intrinsic RasGRP1 gatekeeper of cell proliferation and (b) the extrinsic IL-13 cue, and we postulate that such combination of cell-intrinsic- and extrinsic-cues provide tight control of the tuft lineage while allowing for robust expansion when needed.

The rarity of *Rasgrp1*-expressing cells in the stem/progenitor cell zone in the SI is remarkable, the more because location-wise these cells are distinct from *Lgr5*-positive stem cells. Because of this rarity, *Rasgrp1* transcripts are undetectable in our scRNAseq data set and published datasets and we cannot transcriptionally characterize these cells. Future work will require new tools to analyze the identity of *Rasgrp1*-expressing cells. What is clear is the striking feature of 6,227 DEGs, comparing WT and *Rasgrp1*-deficient TA progenitor cells.

Cell biologically, we demonstrate that RasGRP1 aids to suppress proliferative EGFR signals to license the IL-13 cue. We postulate that such combination of cell-intrinsic- and extrinsic-cues provide tight control of the tuft lineage while allowing for robust expansion when needed. In agreement with a role for modulation of EGFR signals as an intrinsic control mechanism for tuft cell specification, phosphorylation of the EGFR is widespread in these cells (*54*). Our *RoLoRiG* allele, crossed to *Villin^Cre^*, *Villin^CreERT2^*, and *Lgr5^CreERT2^*, and studies combining *in vivo* SI phenotypes with SI organoid findings demonstrate RasGRP1’s impact in TA cells on skewing towards or away from the tuft cell lineage. Decreased numbers of tuft cell numbers have been observed in patients and animal models of inflammatory bowel disease (IBD), while increased tuft cell numbers are reported in patients with irritable bowel syndrome (IBS) (*55, 56*). Together, these findings highlight that a finely tuned equilibrium of tuft cell activity is essential for intestinal health.

Modulation of proliferation in TA cells has been linked to lineage commitment toward absorptive or secretory fate (*5*). DEG analysis of the TA compartment of the WT and *Rasgrp1* deficient SI revealed two remarkable features. First, WT TA cells display broad and varied expression of many transcripts. Second, TA cells without RasGRP1 were present in normal, slightly elevated numbers, were marked as WT counterparts by *Dmbt1* expression, but were completely devoid of this broad and varied transcriptional program. In this body of work, we focused on the trajectory from TA cells to tuft cells, benefiting from the severely reduced *Pou2f3* expression in *Rasgrp1* deficient TA cells and the fact that RasGRP1 perturbations exclusively impact the tuft cell lineage in SI organoids grown in conventional organoid media. In the whole mouse, *Rasgrp1* deficiency or RasGRP1 overexpression via the *Vil^Cre^;R/R* allele impacts other lineages in the SI as well. Future studies, particularly under alternative organoid culture conditions, will be required to determine whether RasGRP1 establishes a poised TA state that broadly facilitates lineage commitment beyond the tuft cell fate.

## Acknowledgements

We would like to thank Christopher Anderson for assistant with initial processing of single-cell RNA sequencing data. We would also like to thank Marine Berquez for technical support with western blot assays. We would like to acknowledge support from the University of California, San Francisco Biological Imaging Development CoLab, and Flow Cytometry CoLab. Finally, we would like to thank past and current members of the Roose and Locksley laboratories for helpful discussions.

## Author contributions

LAS, PD, and JPR conceived the study and the experiments. LAS performed most of the experiments, analyzed the data, and wrote the manuscript. PD and ES initiated the project. Single cell RNA-seq was generated by ES and AS and initial analysis was done by TT. AD and WB helped with 3D organoid lines. HEL infected worms with *Nippostrongylus brasiliensis*. VSC helped with lamina propria isolations. GF, AC and HG supervised scRNA-seq experiments and analysis. CEO and RML provided scientific input. JPR supervised the project and revised the manuscript. All authors read and agreed on the manuscript.

## Data and materials availability

The raw scRNA-seq sequencing data of WT and *Rasgrp1^−/−^*samples have been deposited under GEO accession no. GSE313226. All data are available in the manuscript or supplementary materials.

## Supplementary Materials

**Fig. S1.**
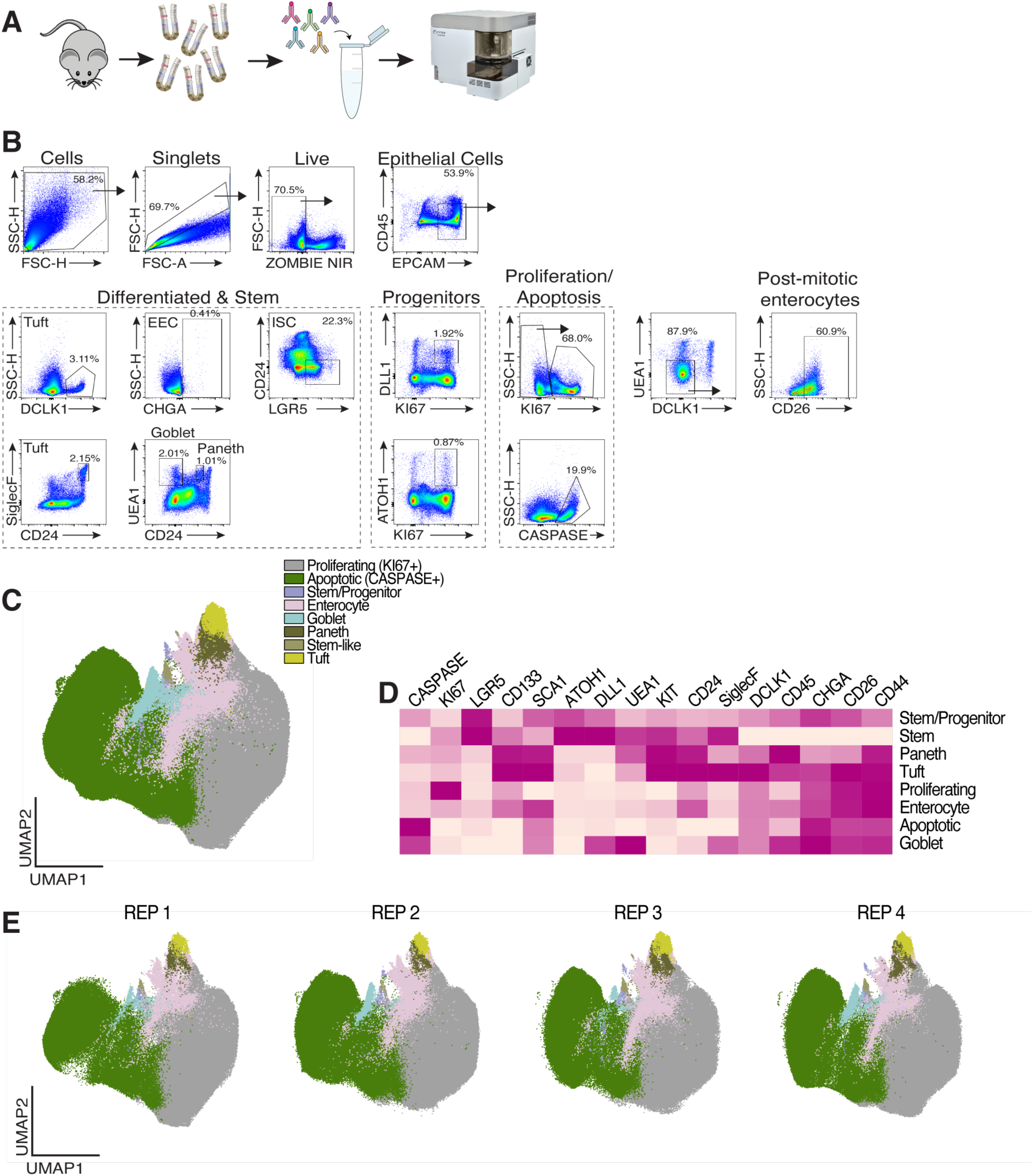
Spectral flow as a tool to assess small intestine cell composition. (**A**) Experimental setup for quantification and analysis of cell types in WT SI using spectral flow cytometry. (**B**) Gating strategy to differentiate intestinal epithelial lineages. (**C**) OMIQ generated UMAP of EPCAM^+^CD45^low/neg^ cells isolated from crypts of 8 C57BL/6 mice, from 4 independent experiments. (**D**) Heatmap of normalized marker expression in each cluster. (**E**) UMAP plots of spectral flow data across four independent experimental days. UMAPs show cells from 2 mice per day, with each panel representing data from a single experimental day. All major cell types are present in each dataset. Data are pooled from 4 separate experiments in C.

**Fig. S2.**
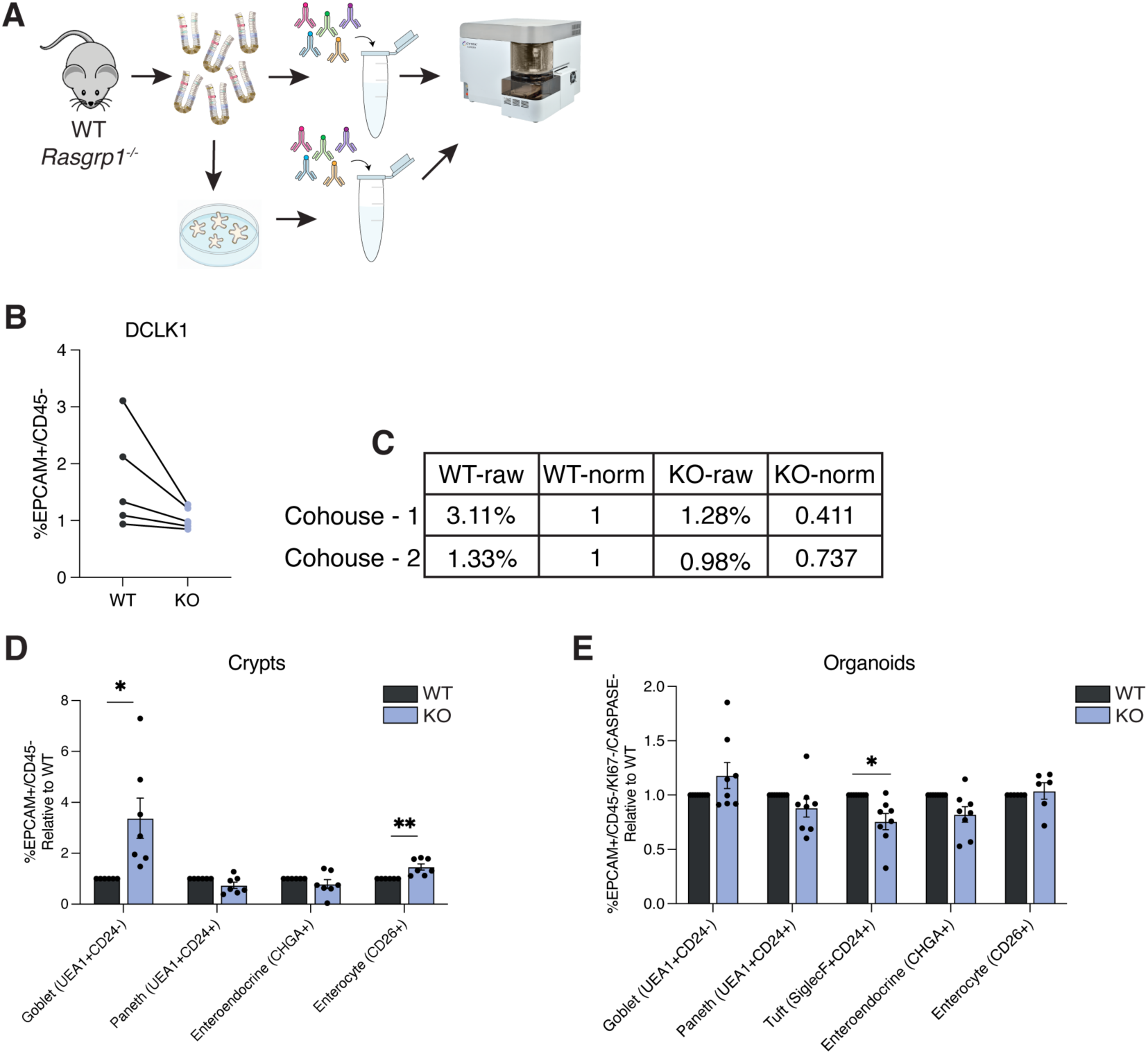
Manual gating analysis of WT and *Rasgrp1^−/−^*intestines and organoids. (**A**) Experimental setup for quantification of cell types in WT and *Rasgrp1^−/−^*SI and organoids using spectral flow cytometry. (**B**) Example of raw spectral flow cytometry data obtained from WT (n=6) and *Rasgrp1−/−*(KO) intestine. Here we used DCLK1 (tuft cell marker) as an example. Mice were cohoused at weaning, and one WT mouse was used as a control for 1-2 KO mice. (**C**) Example of spectral flow batch correction, showing raw versus normalized data from two sets of cohoused WT and KO mice. (**D**) Quantified spectral flow cytometry data of differentiated EPCAM^+^CD45^low/neg^ cells isolated from WT (n=6) and *Rasgrp1^−/−^*(n=7) intestines. (**E**) Quantified spectral flow cytometry data of EPCAM^+^CD45^low/neg^KI67^−^ CASPASE^−^ differentiated cells in WT and *Rasgrp1^−/−^*intestinal organoids. Data in E was normalized as in C, where organoids from each mouse (WT and KO) were cultured in parallel and ran on the same day of culture and passage number. Bars depict mean ± standard error of the mean (SEM). Each data point is a biological replicate (1 mouse, or 2 pooled organoid wells). *p<0.05, **p<0.01 (unpaired two-tailed t-tests). Data are representative of at least three (B, D, E) experiments.

**Fig. S3.**
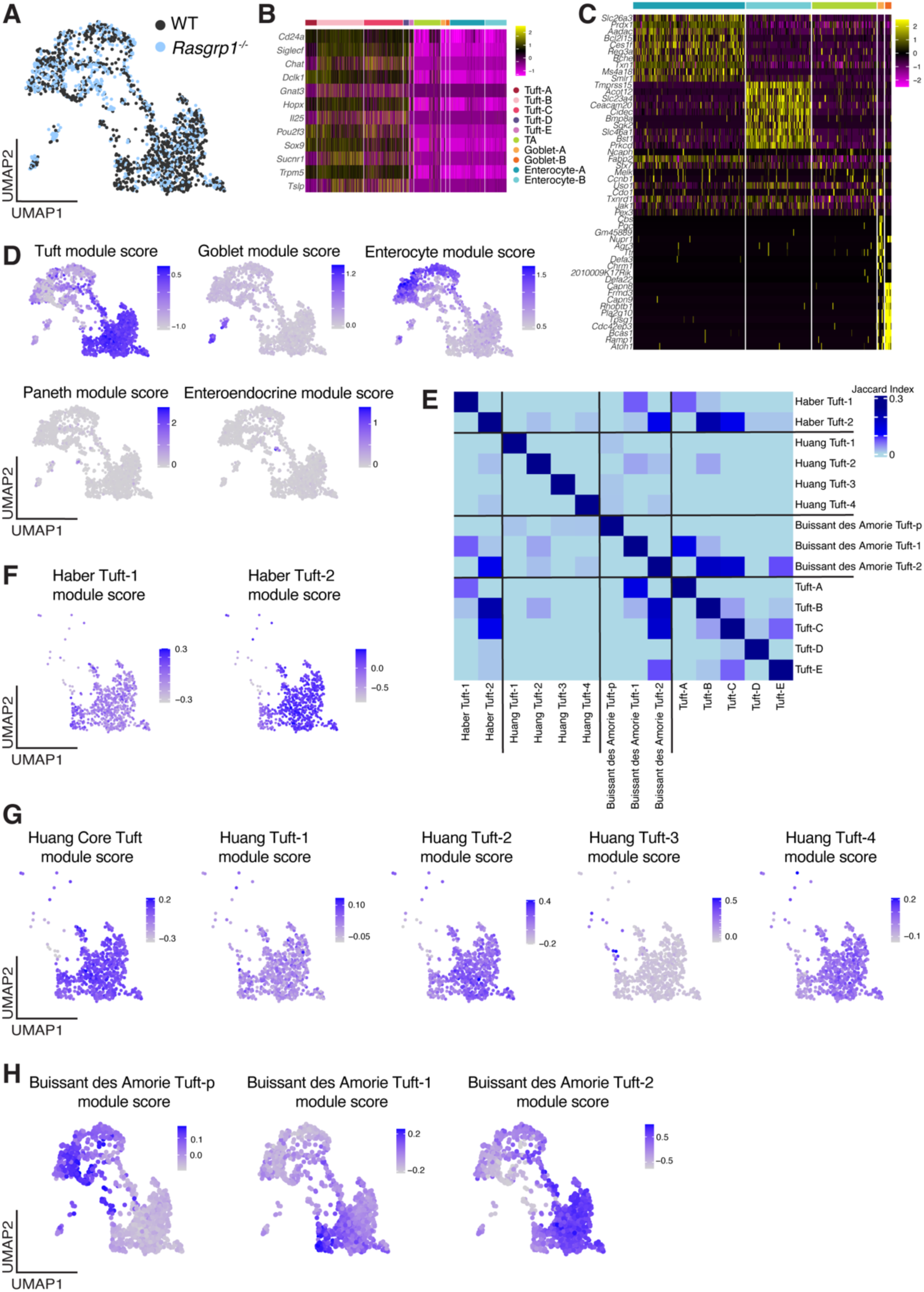
Characterization of tuft cell subsets. (**A**) UMAP visualization of 1,967 EPCAM^+^CD45^neg/low^CD24^+^SiglecF^+^ cells isolated from WT (n=3) and *Rasgrp1^−/−^*(n=3) intestines. Cells are colored by genotype. (**B**) Heatmap of canonical tuft cell genes across clusters in merged dataset. (**C**) Heatmap of the top 20 differentially expressed genes across non-tuft cell clusters in WT dataset. (**D**) UMAPs showing module scores for cell type-specific gene signatures obtained from Haber et al. (Nature, 2017). (**E**) Jaccard similarity index comparing tuft cell clusters from our WT dataset to tuft cell clusters from Haber et al. (Nature, 2017), Huang et al. (Nature, 2024), and Buissant des Amorie (Nature Communications, 2025). (**F, G, H**) UMAPs showing module scores for (**F**) Tuft-1 and Tuft-2 gene signatures obtained from Haber et al. (Nature, 2017), (**G**) core tuft cell genes, Tuft-1, Tuft-2, Tuft-3, and Tuft-4 gene signatures obtained from Huang et al. (Nature, 2024), and (**F**) Tuft-p, Tuft-1, and Tuft-2 gene signatures from Buissant des Amorie et al. (Nature Communications, 2025).

**Fig. S4.**
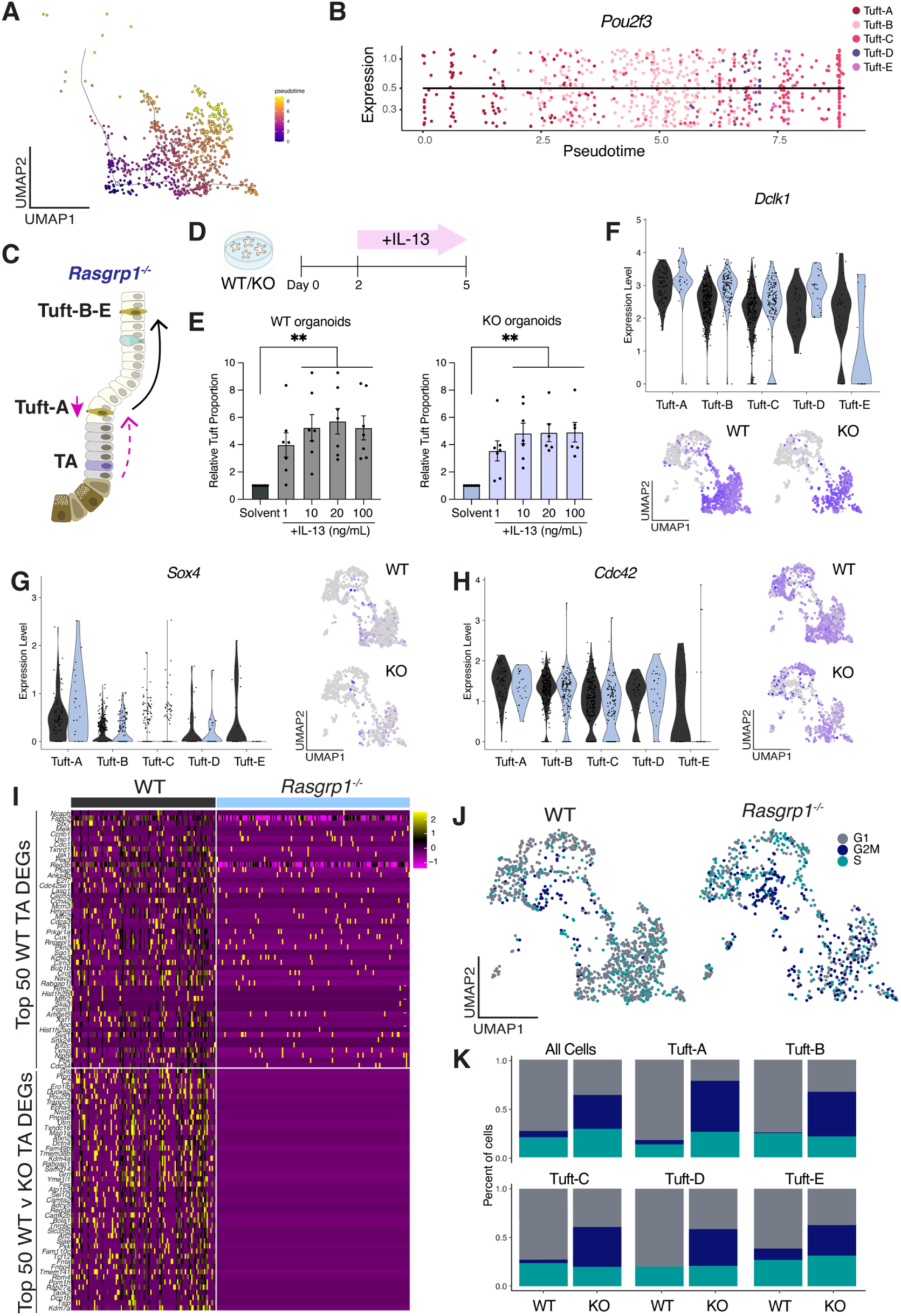
Tuft cell developmental dynamics between WT and *Rasgrp1^−/−^*scRNA-seq datasets. (**A**) UMAP visualization of pseudotime progression of WT tuft cell clusters inferred by trajectory analysis. Pseudotime was computed using Monocle. (**B**) Expression trajectories of *Pou2f3* WT tuft cell clusters over pseudotime. (**C**) Schematic showing effects of *Rasgrp1* deficiency on the tuft cell lineage. (**D**) Experimental diagram for organoid treatments. (**E**) Quantified flow cytometry of DCLK1^+^ tuft cells in WT and *Rasgrp1^−/−^*duodenal organoids treated with increasing concentrations of IL-13. Tuft cell proportions quantified relative to total EPCAM^+^CD45^low/neg^KI67^−^ CASPASE^−^ differentiated epithelial cells. Organoids derived from the same mouse (either WT or KO) were split into control and treatment groups. Treatment values were normalized to the corresponding solvent control from the same passage and treatment day, which was set to 1. Bars depict mean ± standard error of the mean (SEM). Each data point is a biological replicate (2 pooled organoid wells). **p<0.01 (1way ANOVA). (**F-H**) Violin plots and UMAP visualization showing expression of (**F**) *Dclk1,* (**G**) *Sox4*, and (**H**) *Cdc42* in indicated WT and *Rasgrp1^−/−^*clusters. (**I**) Heatmap of the top 50 differentially expressed genes (DEGs) enriched in the WT TA cluster compared with all other clusters (top), and the top 50 genes differentially expressed between WT and KO TA cells (bottom). (**J**) UMAP visualization of WT and *Rasgrp1^−/−^*sorted cells colored by cell cycle phase. (**K**) Percent of WT and *Rasgrp1^−/−^*cells in each phase of the cell cycle per indicated cluster. Data are representative of at least three experiments (E).

**Fig. S5.**
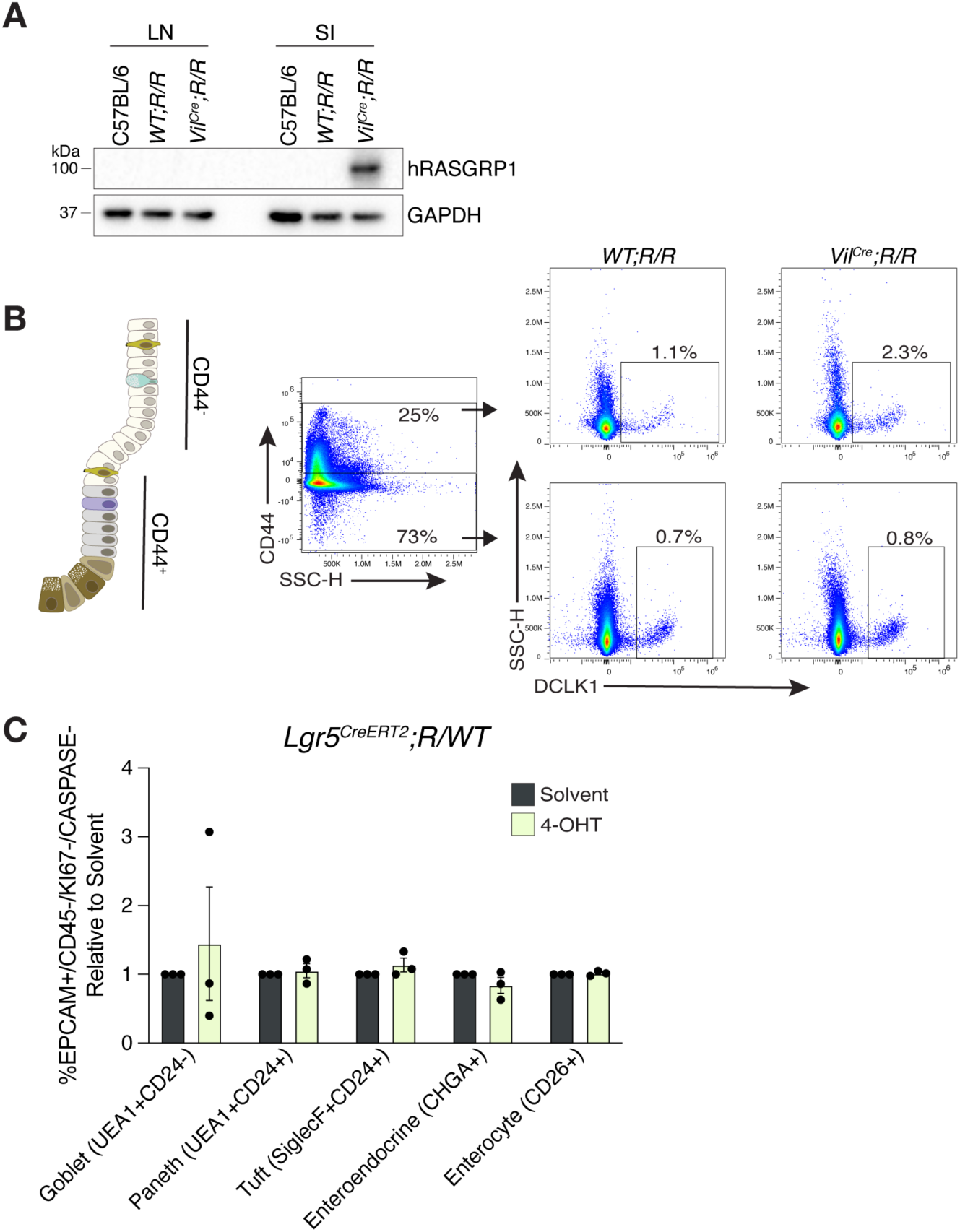
Manual gating and OMIQ analysis of spectral flow cytometry data from *RoLoRIG* intestinal tissue. (**A**) Western blot showing levels of GAPDH and human RASGRP1 in lymph nodes (LN) and small intestine (SI) of C57BL/6, *RoLoRIG;WT* (WT) and *RoLoRIG;Vil^Cre^*(Cre^+^) mice; extended from Figure 4. (**B**) Schematic of CD44 distribution in the SI. Gating strategy and representative flow cytometry plots for quantifying tuft cells in the crypt (CD44^+^) and villi (CD44^−^). (**C**) Quantified spectral flow cytometry data of differentiated EPCAM^+^CD45^low/neg^KI67^−^ CASPASE^−^ cells isolated in 4-OHT and solvent treated *Lgr5^CreERT2^*;*R/WT* organoids. Bars depict mean ± standard error of the mean (SEM). Each data point is a biological replicate (2 pooled organoid wells). Unpaired two-tailed t-tests were run to determine significance. Data are representative of at least two (C) experiments.

**Table S1.** Antibodies used for flow cytometry.

| **Antibody** | **Catalog** | **Company** |
| --- | --- | --- |
| Zombie NIR | 423106 | Biolegend |
| LIVE/DEAD Fixable Blue Dead Cell Stain Kit | L34962 | Invitrogen |
| EpCAM (CD326) | 740559 | BD Biosciences |
| EpCAM (CD326) | 118219 | BioLegend |
| CD45 | M005T02B05 | ThermoFisher |
| CD45 | 103112 | BioLegend |
| CD45 | 109826 | BioLegend |
| Prominin-1 (CD133) | 63-1331-82 | eBioscience |
| Sca-1 (Ly-6A/E) | 746546 | BD Biosciences |
| Sca-1 (Ly-6A/E) | 108124 | BioLegend |
| Delta-like protein 1 (Dll1) | AF3970 | R&D Systems |
| Siglec-F | 155527 | BioLegend |
| Siglec-F | 562068 | BD Biosciences |
| CD24 | 747362 | BD Biosciences |
| CD24 | 101821 | BioLegend |
| c-Kit (CD117) | 105835 | BioLegend |
| c-Kit (CD117) | 105853 | BioLegend |
| CD44 | 612799 | BD Biosciences |
| Ulex europaeus agglutinin (UEA-1) | 73873 | Millipore Sigma |
| Dipeptidyl Peptidase IV (CD26) | 740868 | BD Biosciences |
| Cleaved Caspase3 | 8788S | Cell Signaling |
| Ki67 | 58-5698-82 | Invitrogen |
| Ki67 | 47-5698-82 | ThermoFisher |
| SP-1 Chromogranin A (Bovine) | 20085 | Immunostar |
| DCAMKL1 | ab31704 | Abcam |
| ATOH1 | ab5692 | Millipore Sigma |
| LGR5 | MAB8240 | R&D Systems |
| FlexAble 2.0 CoraLite Plus 647 Labeling Kit | KFA503 | Protein Tech |
| FlexAble 2.0 CoraLite Plus 750 Labeling Kit | KFA504 | Protein Tech |
| Alexa Fluor 700 Conjugation Kit | ab269824 | Abcam |
| Alexa Fluor Pe/Cy5.5 Conjugation Kit | ab102899 | Abcam |

**Table S2. Single cell RNA-sequencing summary, differential expression, and cell cycle phase analysis.**

## Materials and Methods

### Animals

WT C57BL/6 (JAX:000664) and *Rasgrp1^−/−^*mice (JAX:022353) were bred at UCSF. *RoLoRig* mice have been described (*22*), were cross-bred with *Villin-CRE* mice (JAX:004586), *Villin-CreERT2* mice (JAX:020282), and *Lgr5-EGFP-IRES-CreERT2* mice (JAX:008875) and maintained at UCSF. Male and female mice between 7-12 weeks were used in experiments. Mice were housed and treated in compliance with the Institutional Animal Care and Use Committee (IACUC) guidelines at the University of California, San Francisco.

### Histology, Immunostaining, and RNAscope

The duodenum (proximal ∼8-10 cm) was harvested and flushed. For immunofluorescence, tissue was fixed in 4% paraformaldehyde (PFA) for 3 h at 4°C. For RNAscope, tissue was fixed in 4% PFA for 24 h at 4°C. Fixation was followed by a PBS wash and overnight incubation in 30% sucrose. Tissues were coiled into a ‘Swiss roll’ and embedded in Optimal Cutting Temperature Compound (OCT, Tissue-Tek). Samples were stored at −80°C before sectioning (8-12 µm) on a Cryostat (ThermoScientific).

For immunofluorescence, samples were washed with PBS and blocked in blocking solution [Animal free blocking solution (Vector Laboratories, SP-5030), 0.01% SDS, 0.1% Triton X-100] containing 5% normal goat or donkey serum for 1 h at room temperature, then incubated in primary antibody solution overnight at 4°C. The following primary antibodies were used in this study: rabbit anti-DCLK1 (1:1000, Abcam, ab31704), rabbit anti-ECADHERIN (1:200, Cell Signaling, 3195S). Samples were rinsed with PBST (PBS + 0.1% Triton X-100) and incubated in secondary antibody (1:2000) for 1 h at room temperature, followed by DAPI nuclear counterstain (Invitrogen, D1306). Slides were coverslipped with Fluoromount mounting medium (SouthernBiotech, 0100-01).

RNAscope was performed according to manufacturer guidelines using the RNAscope Multiplex Fluorescent Detection Kit v2 (Advanced Cell Diagnostics, 323110). The following probes were used in this study: *Rasgrp1* (486211-C1), *Il13ra1* (437541-C3), and *Sucnr1* (1304611-C2).

Histology images were acquired using a Nikon CSU-W1 SoRa Spinning Disk Microscope and processed using Nikon NIS-elements software and Fiji: ImageJ.

### Intestinal organoids

Murine organoids were generated from crypts isolated from the duodenum (proximal ∼10cm) as described (*14*). Briefly, the duodenum was removed, opened, and cleaned with cold PBS. Tissue was incubated in isolation buffer (PBS, 3mM EDTA) for 30-60 min, rotating at 4°C, then transferred into fresh isolation buffer and manually shaken. Suspension was passed through a 70µm filter to enrich for crypts, which were then plated at ∼500 crypts/well in Matrigel (Corning, 356231) overlayed with complete organoid culture media [Advanced DMEM/F12/Glutamax (Thermo, 10565-018), 10mM HEPES (Thermo, 15630080), 1X Pen/Strep (Life Technologies, 15140-122), 1.25mM N-Acetyl-L-cysteine (Matrix Scientific, 075227), 1X N2 (Life Technologies, 17502-048), 1X B27 (Life Technologies, 12587-010), 100µg/mL Primocin (Invitrogen, ant-pm-1), 50ng/mL murine EGF (Thermo, 53003-018), 100ng/mL murine Noggin (PeptroTech, 250-38), 12.5nM human R-spondin-2 (PeptroTech, 120-43), and 10µM Y-27632 (ATCC, ACS-3030)]. Media was changed every 2-3 days, and organoids were passaged and expanded every 5-7 days at a 1:3 ratio via manual trituration. All experiments were performed on organoids between passages 3-8. For treatment experiments, organoid media were supplemented with 0.5µM or 1µM gefitinib (Selleck Chemicals, S1025), and 1-100ng/mL recombinant IL-13 (BioLegend, 575902) as indicated in figure legends.

For immunostaining, organoids were harvested by incubating on ice in Cell Recovery Solution (Corning, 354253) for 30-40 min, transferred to Eppendorf tubes, and rinsed in PBS/1% BSA. Organoids were fixed in 4% PFA for 30 min at room temperature, washed in PBS, and stored at 4°C in PBS/BSA. Samples were washed with PBS and blocked in blocking solution containing 5% normal goat or donkey serum for 1 h at room temperature, then incubated in primary antibody solution overnight at 4°C. Samples were rinsed with PBST and incubated in secondary antibody for 1 h at room temperature, followed by DAPI nuclear counterstain. Organoids were mounted using Secure-Seal spacers (Invitrogen) stacked onto glass slides to retain structural integrity. Organoid images were acquired using a Nikon CSU-W1 SoRa Spinning Disk Microscope and processed using Nikon NIS-elements software and Fiji: ImageJ.

### Spectral Flow Cytometry

The duodenum was harvested, washed with PBS, and opened longitudinally. For epithelial cell analysis, a crypt-enriched suspension was harvested from the duodenum as described above. Crypts were digested in TrypLE Express supplemented with 10µM Y-27632 and 20µg/mL DNase I at 37°C for 2 min, manually agitating every minute. Cells were washed with FACS Buffer (PBS, 2% FCS, 2mM EDTA, 0.09% Sodium Azide) and passed through a 40µM filter. Lamina propria harvesting and staining was done as described (*24*). Briefly, intestines were incubated with epithelial cell extraction buffer (EEB: HBSS, 3% FCS, 10mM HEPES, 10mM EDTA, 10mM DTT) shaking at 37°C for 15 min. Tubes were vortex at full speed for 30 sec, supernatant discarded, and tissue was placed in fresh EEB to shake for 15 min at 37°C. Tissue was vortexed, supernatant discarded, then placed in rinse buffer (HBSS, 3% FCS, 10mM HEPES) and incubated at 37°C, shaking for 5 min. Tissue was placed in GentleMACS C tubes (Miltenyi Biotec) containing digestion buffer (DB: HBSS, 3% FCS, 10mM HEPES, 30µg/mL DNase I, 0.1 Wunsch/mL Liberase TM), minced with scissors, and incubated with shaking at 37°C for 5 min. To obtain single cells, tissue was subjected to a gentleMACS tissue dissociator (Miltenyi Biotec), using the m_intestine_0 program, filtered through a 100µm filter, then washed with FACS buffer. To harvest organoids, Matrigel was manually dislodged from the plate, and 2 wells were pooled together for dissociation. Organoids were incubated in TrypLE Select for 5 min at 37°C, then transferred to a 96-well v-bottom plate for staining. Cells were washed with FACS buffer then incubated with Zombie NIR (Biolegend) for 15 min at 4°C followed by 2 washes. Cells were next incubated in FACS buffer with antibodies to surface markers (Table S1) for 30 min at 4°C, washed twice, then fixed for 10 min with Foxp3/Transcription Factor Fix/Perm (Tonbo Biosciences). Cells were washed twice with 1X Permeabilization Buffer (Tonbo Biosciences), then incubated with intracellular target antibodies for 45 min at 4°C. Cells were washed twice before acquisition on a 4-laser Cytek Aurora. Spectral flow data were unmixed and compensated using SpectroFlow (Cytek Biosciences, v3.2.1) and further analyzed in FlowJo (version 10.10.0) or uploaded into OMIQ analysis software (https://www.omiq.ai/). For manual gating analysis, cell type proportions were normalized relative to their respective wild-type or littermate control samples. For OMIQ analysis, batch alignment and normalization based on reference samples were performed using CytoNorm. Data were visualized using Uniform Manifold Approximation and Projection (UMAP), and cells were appointed to each cluster using FlowSOM. Debris clusters were removed if they represented less than 0.5% of all cells, or present in only 1 sample.

### Helminth infection

*Nippostrongylus brasiliensis (N. brasiliensis)* infection was initiated in male and female adult C57BL/6, *Rasgrp1^−/−^* and *RoLoRiG* littermates via subcutaneous injection with 500 L3 worms. Mice were euthanized at indicated times and intestinal worm burden was quantified as described (*24*).

### scRNA-seq and analysis

Crypts were harvested from the duodenum (∼10cm) of 3 C57BL/6 (WT) and 3 *Rasgrp1^−/−^*(KO) mice as described (*24*). The intestine was flushed with PBS, opened, and incubated in epithelial cell extraction buffer (Ca^2+^/Mg^2+^ free HBBS, 3% FCS, 10mM HEPES, 10mM EDTA, 10mM DTT) at 37°C with gentle shaking for 15 min. Tubes were vortexed at full speed for 30 sec and tissue was filtered through a 100µm filter. Cells were washed with FACS buffer and incubated with Zombie NIR for 20 min followed by washing. Cells were next incubated with surface markers anti-EpCAM (BioLegend; clone G8.8), anti-SiglecF (BD Biosciences; clone E50-2440), anti-CD45 (Biolegend, clone 30-F11), anti-Ly6C (Biolegend; clone HK1.4), anti-CD24 (BioLegend; clone M1/69), and anti-F4/80 (BD Biosciences, clone T45-2342) for 20 min, followed by washing. Tuft cells were sorted using a FACSAria sorter by gating on CD45^−^ Ly6C^−^ F4/80^−^ EPCAM^+^SiglecF^+^CD24^+^ and collected in tubes containing FCS/50% DMEM. 30,000 cells for each genotype were loaded for 10x Genomics single-cell isolation. Library preparation was performed according to manufacturer’s recommendations. Illumina Hiseq 4000 was used to sequence the sample. Tuft cells of 3 WT and 3 KO mice were pooled per dataset. Sequences were processed with Cell Ranger software by demultiplexing, unique molecular identifier (UMI) collapsing, and alignment to the mm10 mouse transcriptome. The raw data for each dataset generated by Cell Ranger was input into the Seurat R package for preprocessing. Cells were filtered based on quality control metrics: KO cells expressing fewer than 250 genes and WT cells expressing fewer than 500 genes, cells with >25% mitochondrial transcripts, >60% ribosomal transcripts, or more than 100,000 total UMIs were excluded. Filtered counts were log-normalized with a scale factor of 10,000, and highly variable genes were identified. Expression values were scaled, regressing out the effects of total transcript counts, mitochondrial and ribosomal content, as well as S-phase and G2/M-phase cell cycle scores calculated using canonical Seurat gene sets. The WT and KO datasets were then merged into a single object for downstream dimensionality reduction, clustering, and visualization.

Principal component analysis (PCA) was performed on the scaled expression matrix using the top 3,000 highly variable genes. Harmony integration was then applied to correct for potential batch effects between datasets. The first 30 Harmony-corrected principal components were used to construct a shared nearest neighbor graph, which was clustered using the Louvain modularity optimization algorithm. Multiple clustering resolutions were tested, and a resolution of 0.8 was chosen for downstream analysis. Dimensionality reduction for visualization was performed using UMAP on the first 30 Harmony components, and UMAP plots were generated for both the combined dataset and split by library to assess batch correction effectiveness.

Cell clusters were identified based on differentially expressed genes (DEGs) and previously published marker genes. DEGs were determined using a Wilcoxon test through the FindAllMarkers function in Seurat, with the top 50 genes per cluster listed in Table S2. Differential gene expression between genotypes (WT and KO) within each cluster and between cycling and non-cycling cells per cluster was assessed using a Wilcoxon test through the FindMarkers function. Cell type-specific module scores were calculated using the AddModuleScore function on the merged dataset using consensus gene signatures from Haber et al., 2017. For module score analysis on tuft cell clusters, only the WT dataset was analyzed. Module scores were calculated using tuft-1 and tuft-2 genes from Haber et al., 2017, and tuft-1, tuft-2, and tuft-p genes from Buissant des Amories, 2025. Human tuft cell-associated gene programs (tuft-1, tuft-2, tuft-3, tuft-4) from Huang et al., 2024 were converted to mouse orthologs using the gorth function; genes not present in the dataset were excluded prior to module score calculation with AddModuleScore. Pairwise Jaccard similarity between clusters was calculated using the top 50 DEGs per tuft cluster (from Table S2, WT dataset), and corresponding gene lists from published datasets. For each pair of clusters, the Jaccard index was computed as the ratio of the intersection to the union of their respective gene sets. All pairwise indices were assembled into a similarity matrix and normalized using min-max scaling prior to visualization.

Diffusion component analysis was performed using the destiny package (*27, 28*) on WT tuft cell clusters as described (*26*). Tuft cell clusters were subset from the Seurat object, and the first ten principal components were used as input to the DiffusionMap function. Diffusion components 1, 2, and 6 were visualized in three dimensions and used for downstream analysis. Pearson correlation coefficient between diffusion component values and scaled gene expression were calculated to identify the top 50 positively and negatively correlated genes for each diffusion component. The resulting gene sets were analyzed using Metascape (*29*), to determine enriched biological processes and pathway annotations.

Pseudotime trajectory analysis was performed using the Monocle3 package (*38*) on WT tuft cell clusters. Cells were assigned partition information while preserving the original Seurat cluster identities. Cells were then ordered along pseudotime with the tuft-A population designated as the root of the trajectory.

### Immunoblotting

Single cells were harvested from the duodenum (proximal ∼10cm) as described above and lysed with lysis buffer (50 mM HEPES, pH 7.4; 40 mM NaCl; 10 mM sodium pyrophosphate; 10 mM 838 β-pyrophosphate; 50 mM NaF; 2 mM EDTA; 1% Triton X-100) supplemented with protease inhibitor (A32965, Thermo Scientific). Samples were sonicated, centrifuged, and supernatants harvested. Protein content was measured using Pierce BCA Protein Assay Kit (23227, Thermo Scientific). 20 µg of protein was separated on 10% acrylamide gel by SDS-PAGE, then transferred to a PVDF membrane (IPVH00010, EMD Millipore). Membranes were blocked with 5% skim milk in Tris-buffered saline with 0.2% Tween 20 (TBST) then incubated with primary antibody in 5% skim milk in TBST overnight at 4°C. Membranes were washed with TBST and incubated with anti-mouse (PI-2000-1; Vector Laboratories) secondary antibody conjugated with horseradish peroxidase in 5% skim milk for 45 min at room temperature. Membranes were washed with TBST and developed using SuperSignal West Pico PLUS Chemiluminescent Substrate (34580, Thermo Scientific). Images were captured on a ChemiDoc XRS+ system (Bio-Rad).

### Quantitative RT-PCR

Single cells were harvested from the duodenum (proximal ∼10cm) from *N. brasiliensis* infected mice and stained with extracellular antibodies as described above. Cells were sorted directly into RLT buffer (Qiagen) with β-mercaptoethanol using a 5-laser Cytek Aurora. Spectral flow data were unmixed and compensated using SpectroFlow (Cytek Biosciences, v1.4.1). RNA was extracted using a RNeasy Plus Micro Kit (Qiagen), and reverse transcribed with 5X iScript RT Supermix (BioRad). TaqMan Fast Advanced Master Mix (Applied Biosciences) was used for qPCR reactions on a CFX384 Real-Time System (BioRad) machine. Relative gene expression of *Rasgrp1* (4331182; ThermoFisher) was assessed using the ΔΔCT method, with *Actb* (4331182; ThermoFisher) as the housekeeping gene.

### Quantification and statistical analysis

Statistics were calculated by performing unpaired *t* test, one-way analysis of variance (ANOVA) with Tukey’s multiple comparisons post-hoc test, or two-way ANOVA with Sidak’s multiple comparison test using Prism 10 (GraphPad), as indicated in figure legends. Data are mean ± standard error of the mean (SEM) and significance was taken as *p<0.05, **p<0.01, ***p<0.001, and ****p<0.0001.

